# Whole-genome sequencing reveals variant associations with brain imaging phenotypes

**DOI:** 10.64898/2026.01.07.697998

**Authors:** Boxing Liu, Yu Deng, Chao Li

## Abstract

Whole-genome sequencing (WGS) enables comprehensive discovery of genetic variation underlying brain imaging traits beyond the limits of SNP arrays, particularly for rare coding variants with potentially large effects. Here, we integrated WGS data from 45,755 UK Biobank participants with 4,016 derived imaging phenotypes (IDPs) spanning macrostructure (volumes, cortical thickness/area), microstructure (diffusion metrics), functional connectivity, and perfusion MRI measures. We performed variant-level association testing across 2,751,139 coding SNVs under dominant, genotypic, and recessive models. Then, we conducted gene-based rare-variant collapsing analyses across 18,762 genes using multiple qualifying-variant models to improve power for ultra-rare alleles and enhance biological interpretability. We detected 66,893 significant genotype-IDP associations at the variant level and 184 significant gene-IDP associations from collapsing analyses, including many signals driven by protein-truncating and damaging missense variants with larger effect sizes than common variants. Burden heritability analyses supported a contribution of rare coding variation to population variability in IDPs. This study provided a WGS framework for mapping rare coding variant effects on brain structure and function, advancing mechanistic insight into imaging endophenotypes relevant to neuropsychiatric and neurodegenerative disease.

## Introduction

Brain structure and function vary widely across individuals and are closely linked to neurological and psychiatric conditions such as Alzheimer’s disease, schizophrenia, Parkinson’s disease, and autism. Magnetic resonance imaging (MRI) enables the non-invasive quantification of these variations, generating thousands of image-derived phenotypes (IDPs) that serve as informative endophenotypes. These IDPs span macrostructural, microstructural, functional domains, and tissue perfusion, offering a powerful window into the genetic and neurobiological architecture of brain organization, behavior, and complex brain disorders.

Genetic factors are key contributors to these interindividual variations in brain structure and function, motivating large-scale genomic studies to uncover their molecular basis. Recent genome-wide association studies (GWAS) using UK Biobank (UKB) neuroimaging data have identified associations between genetic variants and brain IDPs, particularly for traits such as brain volume, cortical thickness, diffusion parameters, and resting-state functional networks. These efforts have revealed the high heritability and disease relevance of brain IDPs, yet the SNP-array genotyping methods remain largely constrained to common variants, which typically capture population-level heritability but explain only a fraction of total genetic influence and provide limited mechanistic insight. The reliance on SNP arrays further constrains discovery to predefined loci, leaving rare and structural variants underexplored. Consequently, prior imaging-genetics studies^1,2^ have identified that associated loci frequently span large genomic intervals with ambiguous gene attribution. These limitations highlight the need for comprehensive approaches that can bridge common, rare, and structural variants to explain brain phenotypic variation better.

Whole-genome sequencing (WGS) directly addresses the methodological and biological limitations of prior array-based studies by providing complete genome-wide coverage and superior resolution for variant calling, fine mapping, and functional annotation. It captures both rare and common variants across coding, non-coding, and structural regions, allowing for the discovery of previously undetected genetic variation missed by SNP arrays. Rare coding variants often exert larger effects and more directly implicate causal genes, yet have been underexplored due to limited sample sizes and the technological constraints of genotyping and imputation. Large-scale sequencing studies^3–5^ have demonstrated that rare variants contribute independently to phenotypic variance and reveal mechanisms not captured by common-variant GWAS. The development of rare-variant collapsing models enhances statistical power for gene-level discovery, integrating rare and common variation within a unified analytical framework.

Beyond protein-coding variants, regulatory regions, including untranslated regions (UTRs), promoters, enhancers, and long non-coding RNAs (lncRNAs), govern gene expression and neural function, but these regions are poorly captured by SNP arrays or exome sequencing. Evidence shows that variants in such regions can alter transcription initiation, transcript stability, and chromatin architecture, influencing neuronal differentiation, connectivity, and synaptic plasticity^6,7^. Previous imaging-genetics studies focused on protein-coding sequences, leaving the regulatory landscape underexplored. WGS overcomes these gaps by providing base-pair level resolution across the genome and enabling detection of common and rare variants, including novel mutations in non-coding regions. This broader view offers a mechanistic understanding of how genomic variation contributes to brain structure and function.

Structural variants (SVs) and copy number variants (CNVs) form another crucial layer of genomic architecture, complementing single-nucleotide variation. These large rearrangements, including deletions, duplications, inversions, and translocations, can span kilobases, alter gene dosage, and disrupt chromatin topology or long-range regulation, thereby exerting strong effects on brain morphology and disease risk. Array-based neuroimaging studies^8–10^ have shown that CNVs at loci such as 16p11.2, 1q21.1, and 22q11.2 are associated with marked and often reciprocal changes in cortical morphology and white-matter integrity, underscoring gene-dosage sensitivity as a key determinant of neuroanatomical variation and psychiatric vulnerability. However, SNP arrays cannot reliably detect smaller or complex SVs and CNVs, leaving their contribution to brain architecture poorly characterized. WGS fills this critical gap with base-pair precision and improved sensitivity for complex structural variants, enabling integrative analysis of genome organization and its influence on brain structure and function.

Taken together, this study builds on these advances to address the major gaps in prior array-based research: incomplete genome coverage, underrepresentation of coding and non-coding regions, and limited characterization of structural variants. By leveraging WGS data from 45,755 multi-ancestry UKB participants and over 6,000 high-resolution IDPs, we dissect how diverse variants shape brain structure and function. Using WGS-optimized collapsing models, we perform variant– and gene-level association analyses across functional genomic contexts, improving causal resolution and interpretability. We also incorporate SVs and CNVs and conduct ancestry-specific and pan-ancestry analyses to capture the full spectrum of genomic diversity. This integrated design establishes a comprehensive framework for population-scale imaging genetics, advancing mechanistic insight into neuroimaging traits and identifying molecular pathways relevant to neuropsychiatric and neurodegenerative diseases.

## Results

### Overview

We analyzed WGS data from 45,755 UKB participants, integrated with 4,016 high-resolution brain IDPs, to comprehensively map the genetic architecture of brain structure and function. We performed variant-level association tests for all the common and rare variants. We then performed gene-level collapsing analyses, which aggregate the effects of multiple rare variants within genes to improve statistical power and biological interpretability. Building on previously validated rare variant collapsing models, we implemented a multi-model association strategy optimized for WGS, incorporating diverse functional categories including protein-truncating, damaging missense, synonymous, indel, and non-coding variants.

The scanned regions included coding sequences (CDS) and noncoding regulatory segments such as untranslated regions (UTRs), enhancers (single-cell paper), and promoters (definition). In addition, we assessed variants within loci annotated as long non-coding RNAs (lncRNAs), which may regulate gene expression through chromatin remodeling, transcriptional interference, or enhancer-like functions. Furthermore, we tested the annotated structural variations, including CNVs and other large-scale SVs, which can overlap or disrupt both CDS and noncoding regions.

Our primary analysis first focused on individuals of non-Finnish European ancestry (NFE). All association tests applied stringent variant– and gene-level quality control metrics, functional impact annotations, and allele frequency cutoffs tailored to each variant class, ranging from ultra-rare (MAF < 0.002%), rare (MAF < 0.1%) to low-frequency (MAF < 1%) categories.

The brain IDPs spanned macrostructural phenotypes (e.g., global and regional brain volumes, cortical thickness, and area), microstructural measures (e.g., diffusion imaging metrics such as fractional anisotropy and mean diffusivity in white matter tracts), and functional connectivity traits (e.g., independent component analysis (ICA) –based and parcellation-based resting-state network strengths). We further included perfusion metrics, newly derived IDPs obtained from arterial spin labeling (ASL) MRI, capturing cerebral blood flow and volume. These expanded and harmonized phenotypes facilitated the most comprehensive imaging-genetics analysis to date and enhanced the biological interpretability of discovered loci.

### SNV variant-level associations

*<u>CDS</u>*: We performed a variant-level association study to test for associations between all 4,016 phenotypes and 2,751,139 protein-coding region SNVs observed in at least six participants of European ancestry (i.e., MAF>0.0006492%). We defined three genetic models: dominant, genotypic, and recessive(see Methods), equating to 33,517,126,437 tests. We used a two-sided Fisher’s exact test for binary traits and linear regression for quantitative traits. Using a *P-*value threshold of P ≤ 1×10−8 (Figure 1), and excluding the MHC region (chromosome 6: 25–35Mb), we identified 66,893 genotype-IDP associations for quantitative traits (dominant model: 50,464, genotypic model: 67,918, recessive model: 20,747) (Supplementary Table XXX) covering 2,185 common genotype signals (MAF > 1%), 32 low-frequency genotype signals (MAF: 0.1%-1%), 165 rare genotype signals (MAF: 0.001%-0.1%) and 54 ultra-rare genotype signals (MAF < 0.001%).

**Figure 1:**
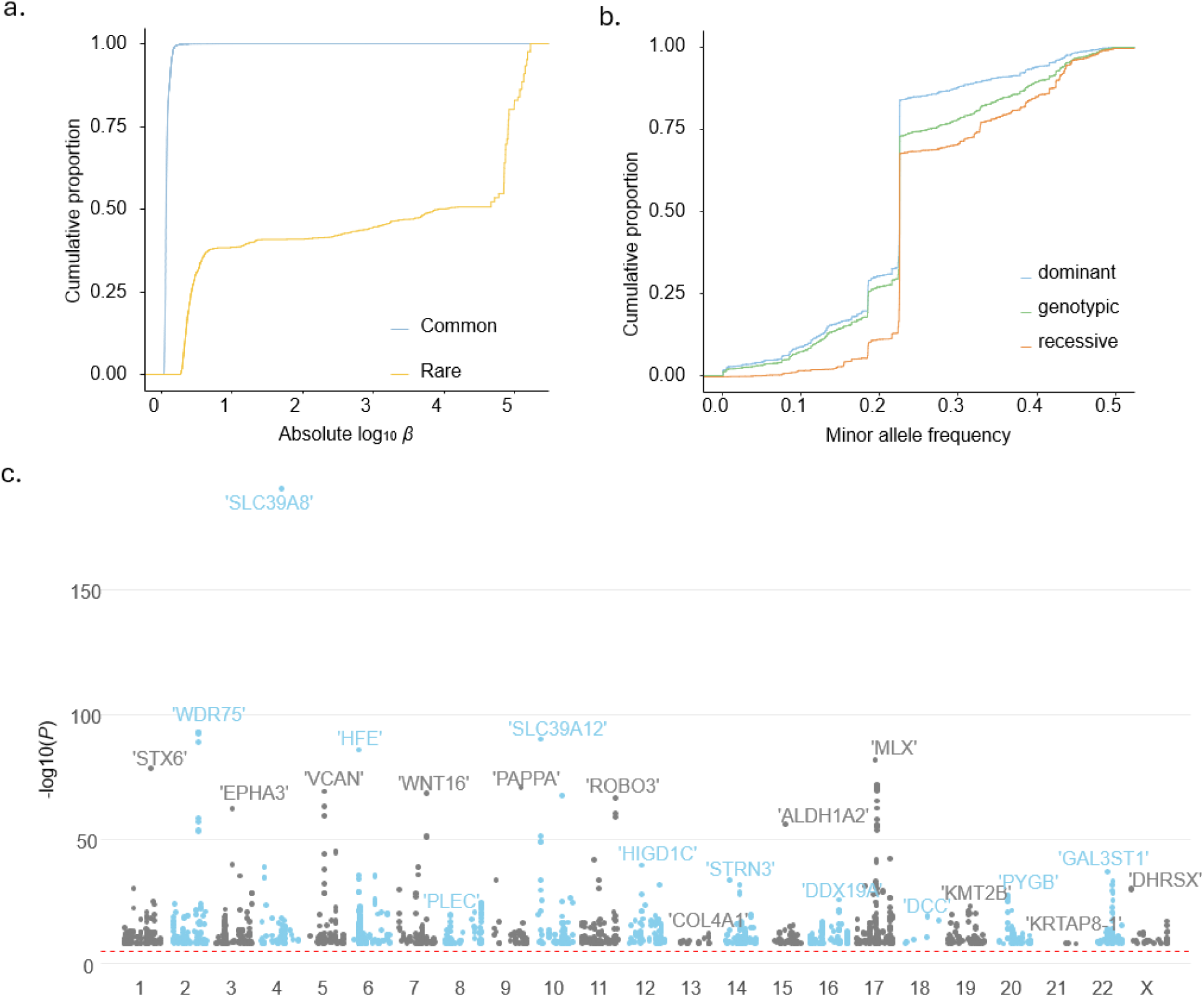
a. The distribution of effect sizes for common versus rare significant (p < 1e-8) variants in CDS. b. The MAF distribution of genome-wide significant (p < 1e-8) variants associated with imaging-derived phenotypes in CDS. c. Genomic distribution and significance of the associations (p < 1e-8) in CDS. The labeled genes represent the most significant gene within each locus.

At the rarest significant MAF (cohort MAF of 0.0013%), we identified 13 significant signals. In particular, two significant variants were a frameshift variant and a stop-gained variant in calcitonin receptor CALCR and FAT atypical cadherin 2 (*FAT2*), respectively. *CALCR* was associated with CSF normalised volume, global total grey matter volume, MD, and L3 value of the left superior fronto-occipital fasciculus, L2 and L3 value of the left fornix cres with stria terminalis (Supplementary Table 3). *FAT2* was associated with global total grey matter volume, right cerebellum cortex volume, and left cortex volume.

Common variants replicated many previously identified GWAS signals while refining the localization of causal regions. The distribution of effect sizes followed the expected inverse relationship with allele frequency, and rare variant associations explained additional trait variance beyond common variants.

The effect sizes of significant rare variant associations were substantially larger than those of common variants (Fig. 1a). While some significant variants are probably in linkage with nearby causal variants, associated PTVs and missense variants often represent the causal variant themselves, implicating protein-coding genes involved in neurodevelopmental and synaptic pathways.

### Rare variant collapsing analyses

We also performed gene-level association tests using collapsing analyses. In this approach, the proportion of cases with a qualifying variant was compared with the proportion of controls with a qualifying variant in each gene. We used 12 different sets of qualifying variant filters (models) to test the association between 18,762 genes and 4,061 phenotypes (Methods), equating to 76,192,482 tests. The models included ten dominant models, one recessive model, and one synonymous variant model that served as an empirical negative control (Methods).

Gene-level collapsing analyses identified 184 significant gene-IDP associations (P ≤ 1×10−8), complementing the variant-level discoveries (Fig. 2). Many signals were of large effect, with a median absolute beta of 1.04 for quantitative traits.

**Figure 2:**
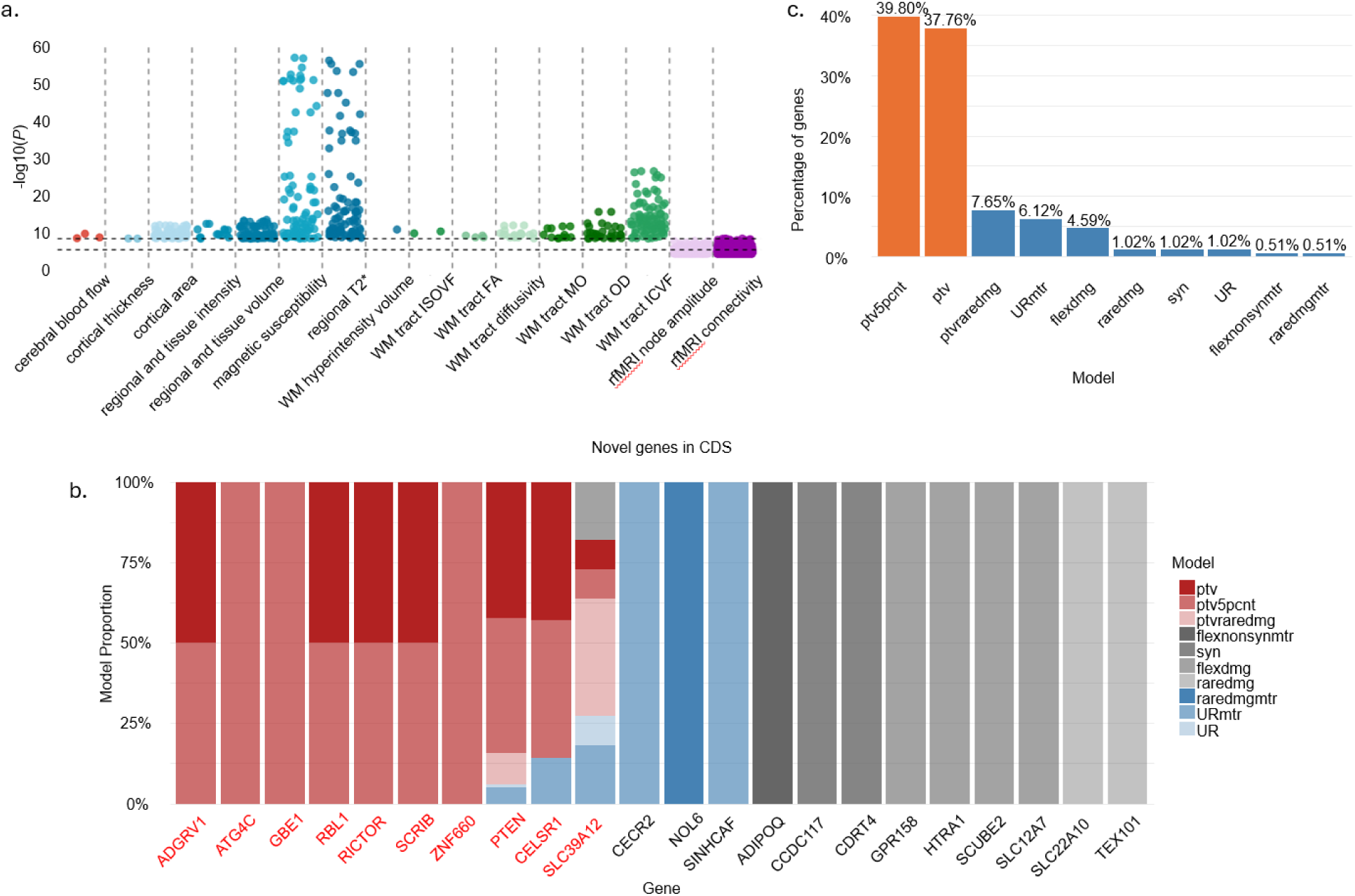
a. Gene-level association results of imaging categories from whole-genome sequencing (WGS) across CDS. Each point represents a gene with P < 1e-5, and the dashed horizontal line marks the threshold (P < 1e-8). b. Novel significant genes identified in the coding sequence (CDS) region. The y-axis shows the proportion of different genetic models contributing to the association signal. Genes labeled in red were identified through protein-truncating related variants (PTVs). c. The proportion of novel genome-wide significant genes identified by each genetic model. The top two models, ptv5pcnt (39.80%) and ptv (37.76%), account for the majority of identified genes, indicating a predominant role of protein-truncating variants (PTVs) in driving significant associations.

Collapsing models focused on PTVs explained 52% of quantitative associations. Remaining signals emerged from models that included missense variants. While these results confirm the importance of PTVs, they also emphasize the role of other forms of variation in human disease.

Overall, these genes cluster into several major functional categories: iron and metal ion transport and homeostasis represented by *TF*, *SLC11A2* and *SLC39A12*; immune–vascular and scavenger-receptor pathways and cell-adhesion processes represented by *STAB1*, *SCUBE2*, *ADGRV1* and *CELSR1*; glucose–lipid metabolism and energy homeostasis represented by *ADIPOQ*, *GBE1* and *LRP5*; and the PI3K–AKT–mTOR and cell-growth signaling axis represented by *PTEN*, *RICTOR*, *PLCB4*, *PLEKHG3,* and *RBL1*. In addition, genes such as TUBB3 and SCRIB are closely involved in neurodevelopment and cytoskeletal remodeling. Together, these findings indicate that the significant PTV-bearing genes identified in our study are not randomly distributed across the genome but are enriched in a limited number of biological pathways related to metabolism, immunity, and cell-growth regulation, thereby providing important clues to the shared molecular mechanisms underlying diverse complex traits.

We identified a set of highly pleiotropic genes that showed significant effects across multiple traits, with *TF* (146 gene-IDPs associations), *STAB1* (123 gene-IDPs associations), and *PTEN* (118 gene-IDPs associations) displaying the largest numbers of associated phenotypes, suggesting that they occupy key positions within the underlying biological networks. Specifically, we identified 29 associations between *TF* and T2* in the caudate nucleus, putamen, pallidum, thalamus, and substantia nigra. Among these results, associations with the caudate nucleus, putamen, and pallidum were consistent with Elliott et al in GWAS analysis^1^. Moreover, we also identified 25 associations between *TF* and magnetic susceptibility-related IDPs in the same anatomical structures with T2*, including the hippocampus. In addition, 78 associations between *TF* and ICVF of white matter tracts were identified, including cerebral peduncle, anterior limb of internal capsule, superior cerebellar peduncle, posterior limb of internal capsule, fornix cres & stria terminalis, and medial lemniscus. Consistent with TF, we also identified *SLC11A2* and *SLC39A12* associated with magnetic susceptibility-related IDPs and T2*. All genes are known to affect iron transport and storage, or neurodegeneration with brain iron accumulation (NBIA). In addition, 46% (66/146) of associations were identified by PTV models.

*STAB1* encodes the scavenger receptor Stabilin-1, which is predominantly expressed in vascular endothelial and immune cells and is involved in ligand clearance and immune regulation. The largest number of associations were magnetic susceptibility-related IDPs (51 associations) and T2* (52 associations) as well, including caudate nucleus, putamen, pallidum, accumbens, thalamus, and substantia nigra, with 90% (111/123) PTV models. It is noted that *TF* was negatively associated with T2* and positively associated with median magnetic susceptibility, while *STAB1* was reversedly.

PTEN is a classical tumor suppressor gene encoding a lipid phosphatase that constrains cell growth by inhibiting the PI3K–AKT signaling pathway. The largest number of associations were cortical area (61 associations) and regional and tissue volume (55 associations), covering vital brain regions, such as total brain surface area, volume of cerebral white matter, grey matter, and amygdala. In particular, 94% (111/118) of signals were identified by PTV models.

We identified 22 novel IDP-associated genes, as shown in Fig. 2b, including *ADGRV1, GPR158, SLC39A12, CELSR1,* and *HTRA1*, which are all involved in neurodevelopment, synaptic function, or cerebral small vessel disease. These collapsing results demonstrate that aggregating rare variants enhances detection power for biologically interpretable signals, particularly when individual variants are too infrequent to reach genome-wide significance. Moreover, gene-level hits highlight convergence between rare and common variant architectures within shared pathways, emphasizing the complementary roles of different variant classes in shaping brain structure and function.

### Heritability analysis and genetic correlation

We first estimated the heritability of brain IDPs attributable to rare variants using burden heritability regression (BHR). Following BHR best practices, variants were stratified into three frequency bins: low-frequency (0.01-0.001), rare (0.001-1e-5), and ultra-rare (<1e-5) and four functional impact categories as defined by the annotation tool (package ‘SnpEff’): High (e.g., protein truncation, loss of function, or nonsense-mediated decay), Moderate (e.g., missense variants that alter amino acids), Low (typically synonymous or minimal impact), and Modifier (variants located in non-coding or regulatory regions with less predictable effects).

Among ultra-rare variants, high-impact variants exhibited the highest heritability (median h² = 0.0047, max h² = 0.014), followed by modifier variants (median h² = 0.0076, max h² = 0.0157). moderate (median h² = 0.0020) and low (median h² = 0.00095) categories showed lower heritability. For rare variants, Moderate impact variants contributed the most heritability (median h² = 0.00798), followed closely by low (median h² = 0.00630), with high (median h² = 0.00541) and modifier (median h² = 0.00231) variants showing smaller effects. In the low-frequency variants, low-impact variants dominated (median h² = 0.01346), followed by moderate (median h² = 0.01256), with minimal contributions from high and modifier variants (Fig. 3).

**Figure 3:**
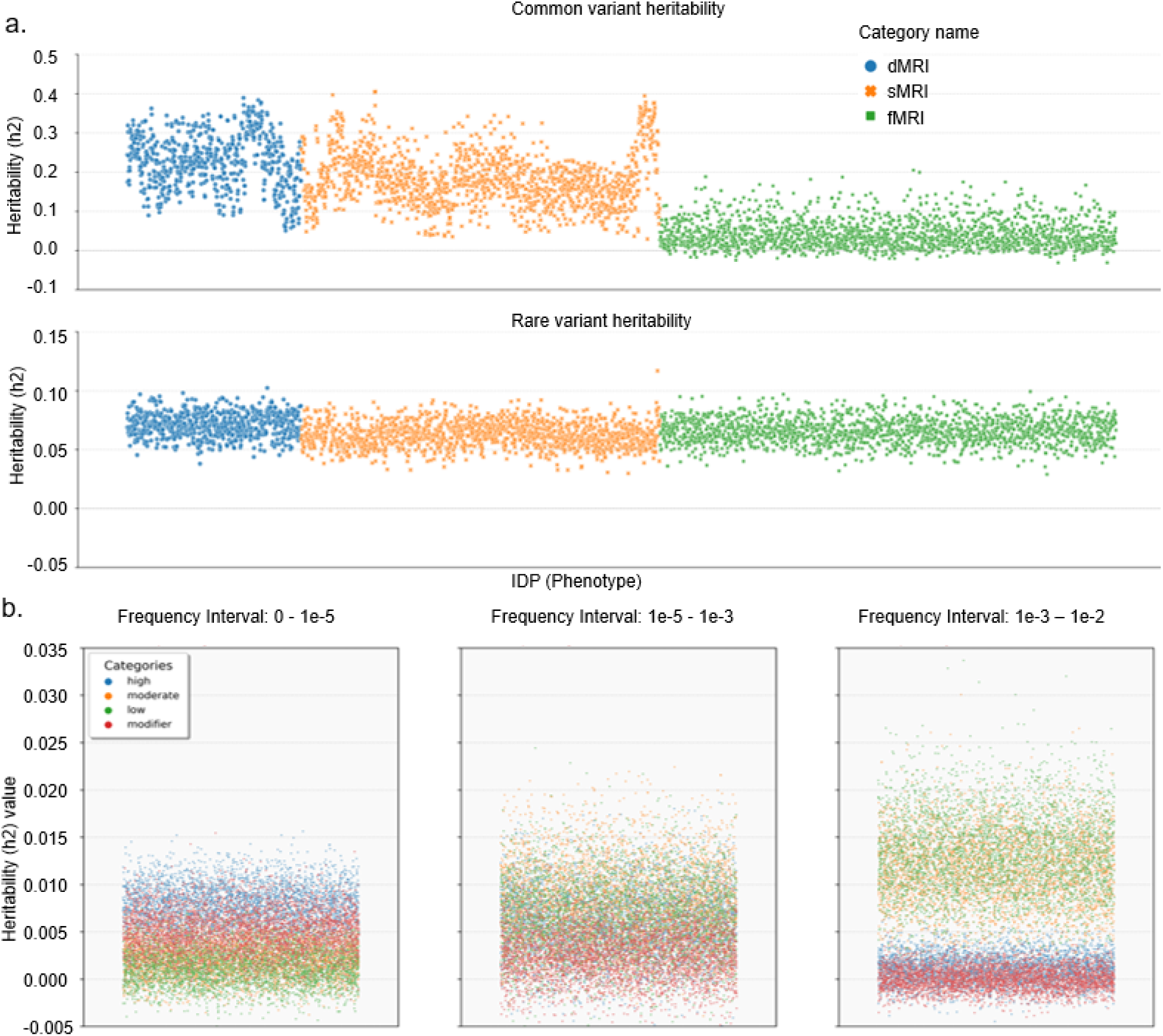
a. Top: SNP-heritability (h²) explained by common variants for each imaging-derived phenotype (IDP). Bottom: SNP-heritability (h²) explained by rare variants for the same set of IDPs. Each point represents one IDP and is coloured by imaging modality: blue, dMRI; orange, sMRI; green, fMRI. b. Left, middle, and right panels correspond to different MAF bins: ultra-rare (MAF < 1×10⁻⁵), rare (1×10⁻⁵-1×10⁻³), and low-frequency (1×10⁻³–1×10⁻²), respectively. Points are coloured by SnpEff functional category: blue, High; orange, Moderate; green, Low; red, Modifier.

Our results demonstrate that the contribution of rare variants to brain imaging phenotypes varies not only by allele frequency but also by predicted functional impact. Notably, High-impact variants showed the strongest heritability within ultra-rare alleles, consistent with their severe functional effects. At higher frequency bins, particularly in the rare and low-frequency ranges, moderate– and low-impact variants dominated, reflecting the greater statistical power to detect weaker but more prevalent effects.

Aggregated across categories, total burden heritability for all IDPs ranged from 0.05 to 0.1 (median h² = 0.0653, max = 0.1170). Among IDP categories, the T2* signal showed the highest median heritability (0.0771), while cortical gray-white contrast exhibited the lowest (0.0567). For comparison, common variant heritability in previous GWAS studies explained approximately 5-10% of variance across many IDPs, with peak estimates up to 0.35 for global and regional brain volumes. Our findings show that rare variants, especially those ultra-rare and of high functional impact, can explain a comparable or even greater proportion of variance for certain traits, highlighting that the contributions from sparse, high-impact variants are significant, comparable to polygenic common variant effects. Importantly, trait-wise concordance between rare and common variant heritability patterns suggests their complementary roles in shaping brain structure and function.

Using BHR, we next computed burden-based genetic correlations across cortical traits, including volume, surface area, gray-white contrast, and thickness, based on the DKT atlas parcellation. We observed significant correlations across all features, indicating shared genetic underpinnings. Gray-white contrast exhibited the strongest median genetic correlation (|r| = 0.4953), followed by cortical thickness and surface area, while volume showed the weakest correlation (|r| = 0.2730), suggesting trait-specific genetic influences.

Further analysis within cortical lobes revealed that intra-lobe genetic correlations were consistently higher than inter-lobe correlations. Notably, the parietal lobe showed the highest internal consistency across features, with |r| = 0.6184 (area), 0.5783 (thickness), 0.7543 (gray-white contrast), and 0.4849 (volume). This pattern suggests that the genetic determinants of structural features may exhibit regionally specific clustering.

While previous GWAS based on common variants^1,2^ have shown similar patterns of regional and trait-specific heritability, our study demonstrates that rare variant burden also recapitulates this spatial architecture. Importantly, we find that the strongest genetic correlations in rare variant models occur for traits that previously showed high SNP heritability, such as gray-white contrast and cortical area. This concordance suggests that both common and rare variants act on overlapping biological substrates, but rare variants, particularly those with high predicted functional impact, may offer additional mechanistic insights. Hence, our findings highlight the complementarity of rare variant burden analyses in enriching the neurobiological interpretation of imaging-genetic architecture.

### Conditional collapsing analysis

To assess whether rare variant associations were independent of nearby common variant effects, we performed conditional gene-based collapsing analyses. For each significant gene, GWAS were conducted using imputed genotype data from UKB v3 (PLINK v2.0), focusing on common variants (MAF > 0.5%) within ±500 kb of each gene boundary. Significant common variants (P < 1 × 10⁻⁵, r² < 0.01 after clumping) were then incorporated as covariates in re-run collapsing models to control for local LD effects. This conditional approach allowed us to disentangle the contributions of rare and common variants to gene-level signals. Across all significant genes, 80% of associations remained significant after conditioning, suggesting that most rare variant associations represent independent effects rather than tagging of common variant loci. Notably, independent rare variant signals were particularly enriched in genes involved in (insert pathway examples, e.g., axonal transport, ion channel regulation), whereas shared loci often mapped to well-known GWAS hits. These results provide evidence for complementary architectures: rare variants with large, independent effects alongside common variants contributing polygenically within shared biological pathways.

### Biological annotation of the identified genes

To assess the biological relevance of genes identified through both single-variant and gene-level rare variant association tests, we performed pathway enrichment and tissue-specific expression. Pathway and tissue enrichment analyses demonstrated that genes identified across variant classes converge on shared biological processes, including synaptic signaling, axon guidance, neuronal morphogenesis, and glial regulation.

#### Tissue Enrichment

Across all trait groups, GTEx-based tissue enrichment consistently pointed to brain tissues. The top enriched tissues included the amygdala, hippocampus, putamen, anterior cingulate cortex, and frontal cortex, regions central to emotion, cognition, and memory (Fig. 4). This pattern overlaps substantially with common variant studies, where trait-associated loci were also enriched for brain-expressed genes with the highest specificity in cortical and subcortical areas^1,2^, supporting convergence of common and rare variant effects on shared neuroanatomical circuits.

**Figure 4:**
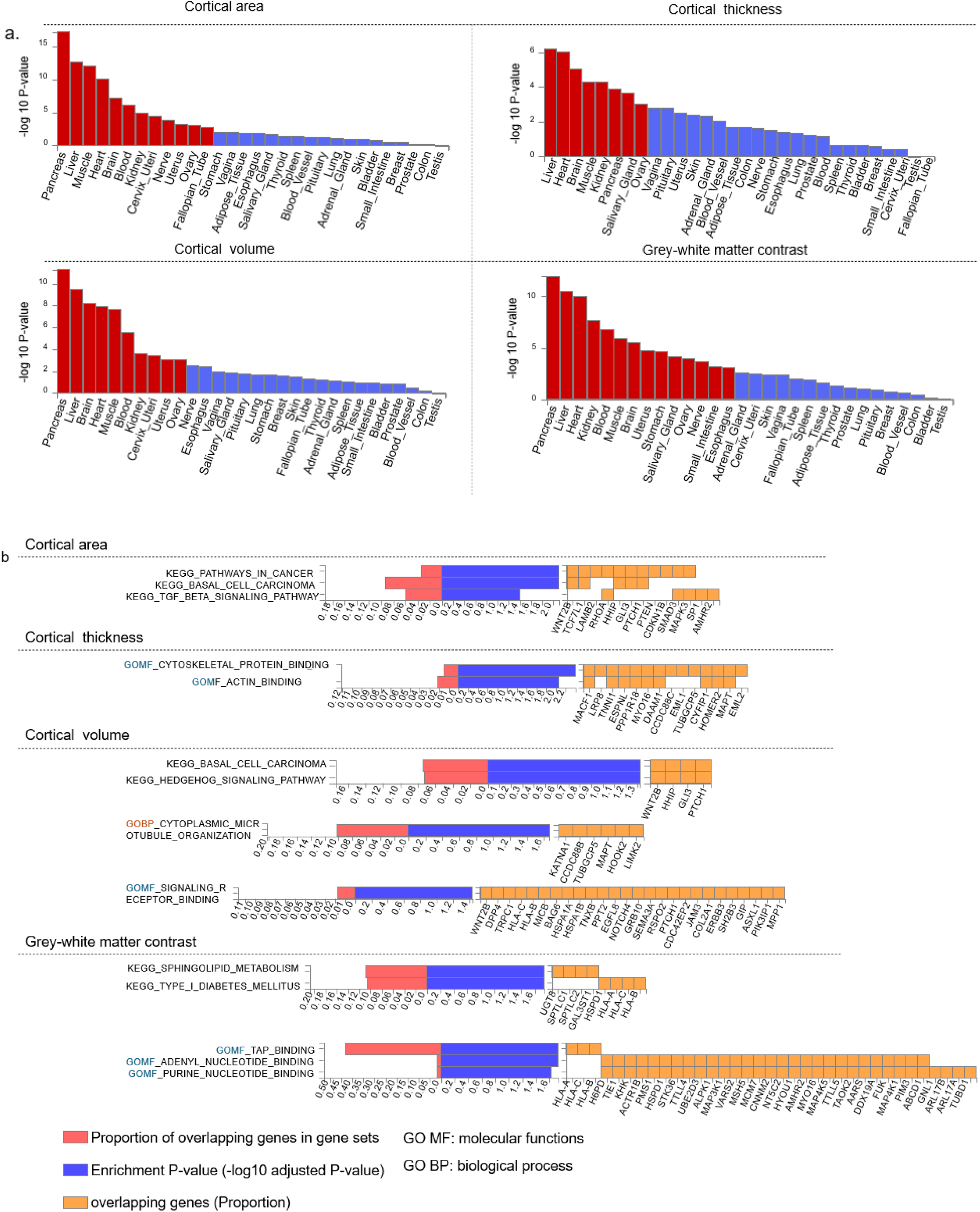
a. Differential expression of brain IDP-associated genes across GTEx tissues. Bar plots show −log₁₀ P values for enrichment of differentially expressed gene (DEG) sets in each tissue for four cortical phenotypes: cortical area (top left), cortical thickness (top right), cortical volume (bottom left), and grey–white matter contrast (bottom right). Tissues are ordered by enrichment strength along the x-axis. Bars highlighted in red indicate DEG sets that remain significantly enriched after Bonferroni correction (P < 0.05); non-significant tissues are shown in blue. b. Pathway enrichment of cortical trait–associated genes in KEGG and Gene Ontology (GO). For each phenotype (cortical area, cortical thickness, cortical volume, and grey–white matter contrast), the most significantly enriched KEGG, GO molecular function (MF), and GO biological process (BP) pathways are shown. Red bars indicate the proportion of input genes overlapping each pathway, blue bars show the –log₁₀ adjusted enrichment P value, and orange boxes list the overlapping genes (with their proportion indicated on the x-axis).

#### Pathway enrichment

For cortical areas, KEGG pathway analysis revealed significant enrichment in Pathways in Cancer (P = 7.76×10-5) and Basal Cell Carcinoma (P = 8.38×10-5), suggesting pleiotropic roles in proliferative and morphogenetic pathways relevant to neurodevelopment. Gene Ontology (GO) analysis further highlighted regulatory processes such as response to stimulus (P = 1.36×10-5), protein polymerization (P = 2.09×10-5), and organelle organization (P = 2.94×10-5), indicating cytoskeletal remodeling and intracellular transport as potential mechanisms influencing cortical expansion (Figure 4). These findings align with prior GWAS reports of cortical areas implicated in cell growth and adhesion processes, now extended to rare variants with putatively larger effect sizes.

For gray-white matter contrast, Enriched pathways include sphingolipid metabolism (P = 1.76×10-4) and diabetes mellitus (P = 1.96×10-4), as well as binding-related GO terms (TAP binding, adenyl nucleotide binding, purine nucleotide binding; P < 4×10-5), suggesting a mechanistic link between membrane composition, myelination, and axonal signaling. These processes have also been highlighted in common variant studies associating lipid metabolism genes with white matter microstructure^11^, suggesting converging biological pathways across variant classes.

For cortical thickness, GO terms including actin binding (P = 9.81×10-6) and cytoskeletal protein binding (P = 2.25×10-6) reflect molecular determinants of cortical laminar architecture, echoing previous findings where cytoskeletal genes (e.g., DAAM1, TUBB) were linked to regional thickness in common variant GWAS^1^. Our results support a similar biology driven by rare coding variants, potentially affecting neurogenesis and cortical stability.

For cortical volume, Enriched pathways included Basal Cell Carcinoma (P = 4.75×10-4) and signaling receptor binding (P = 2.1×10-5), suggesting roles in mitogenic and extracellular signaling. This is consistent with previous SNP-based studies where brain volume was associated with general growth factor signaling^2^.

## Discussion

We integrated whole-genome sequencing data from 45,755 UK Biobank participants of non-Finnish European ancestry with 4,016 MRI-derived phenotypes, focusing exclusively on coding sequences to map the contribution of protein-altering variants to brain structure and function. Across 2,751,139 coding SNVs observed in at least six individuals, we identified 66,893 significant variant–IDP associations and 184 significant gene-IDP associations from gene-level collapsing analyses. Many signals overlapped or refined loci previously reported in array-based imaging GWAS, whereas others were novel or extended known biology to additional cortical, subcortical, and white-matter traits. Rare and ultra-rare protein-truncating and damaging missense variants, including frameshift and stop-gained changes in genes such as *CALCR* and *FAT2*, showed effect sizes that far exceeded those of common variants and were associated with global and regional volumes, diffusion metrics, and susceptibility-based measures.

Gene-level collapsing models, which aggregated private-to-rare functional variants, captured many associations that were not detectable in single-variant tests and revealed convergence on a restricted set of pathways. These included iron and metal ion homeostasis (*TF*, *SLC11A2*, *SLC39A12*), immune–vascular and scavenger-receptor signalling (*STAB1*, *SCUBE2*, *ADGRV1*, *CELSR1*), glucose–lipid metabolism and energy homeostasis (*ADIPOQ, GBE1, LRP5*), and PI3K–AKT–mTOR and cell-growth regulation (*PTEN, RICTOR, RBL1*, with *TUBB3* and *SCRIB* implicated in neurodevelopment and cytoskeletal dynamics). Highly pleiotropic genes such as *TF, STAB1, and PTEN* showed extensive effects across T2*, susceptibility, and cortical morphometry, suggesting that modest dosage perturbations in central regulators of metal handling, vascular–immune function, and growth-factor signalling can influence multiple neuroanatomical systems. Notably, we identified coding variants in *ADGRV1, GPR158, SLC39A12, CELSR1, and HTRA1* that are all involved in neurodevelopment, synaptic function, or cerebral small vessel disease, supporting the idea that mechanisms underlying rare monogenic disorders also modulate subclinical population-level variation in brain structure.

Our variant-level analyses used Fisher’s exact tests for binary traits and linear regression for quantitative traits, following previously validated rare-variant association frameworks, whereas gene-level collapsing models were optimized for protein-truncating and damaging missense variants and benchmarked against a synonymous negative-control model to define a study-wide significance threshold. As in other sequencing studies, Fisher’s exact tests cannot adjust for covariates and therefore require stringent quality control, careful case–control harmonization and ancestry pruning; in the absence of such measures, regression-based approaches with explicit covariate adjustment will be essential, and future work should systematically benchmark these methods for imaging phenotypes. Burden heritability regression restricted to coding variants showed that rare and ultra-rare protein-altering variants explain a non-trivial proportion of variance in many IDPs and that their burden-based genetic correlations across cortical traits broadly mirror SNP-based patterns, indicating that rare and common coding variants act on overlapping biological substrates.

Our analyses are specific to European ancestry, and we are underpowered to detect many ancestry-specific signals and do not capture non-coding or structural variation. Larger, more diverse WGS-MRI cohorts and integration with regulatory and single-cell data will be required to refine the mechanisms proposed here and to fully delineate the contribution of non-coding and structural variants. Overall, these results demonstrate that rare and ultra-rare coding variants constitute an important, mechanistically interpretable component of the genetic architecture of brain imaging traits and provide a gene-resolved framework for linking protein-coding variation to neuroanatomical organization and risk for neuropsychiatric and neurodegenerative disease.

## Methods

### Imaging-derived phenotypes (IDPs)

We analysed brain IDPs from the UKB “40k” imaging data release (around 40,000 participants) from early 2020, processed by WIN FMRIB on behalf of UK Biobank^12^. After excluding individuals during genetic quality control (see below), the final sample comprised 45,755 participants.

As detailed in ref.^13^, the UKB imaging protocol includes six MRI modalities: T1-weighted and T2-weighted fluid-attenuated inversion recovery (T2-FLAIR) structural scans, susceptibility-weighted imaging, diffusion MRI, task-based fMRI, and resting-state fMRI. We further included perfusion metrics, newly derived IDPs obtained from arterial spin labeling (ASL) MRI, capturing cerebral blood flow and volume, and quantitative magnetic susceptibility^14^. Totally, we obtained 4,016 IDPs, and grouped the IDPs into 19 categories, including arterial transit time, cerebral blood flow, cortical area, cortical grey-white contrast, cortical thickness, magnetic susceptibility, regional T2*, white matter hyperintensity volume, rfMRI connectivity, rfMRI node amplitude, tfMRI activation, white matter (WM) tract FA, WM tract diffusivity, WM tract ICVF, WM tract ISOVF, WM tract MO, and WM tract OD.

### WGS processing, QC, and variant calling

WGS data of the UKB participants were generated by deCODE Genetics and the Wellcome Sanger Institute as part of a public-private partnership involving AstraZeneca, Amgen, GlaxoSmithKline, Johnson & Johnson, Wellcome Trust Sanger, UK Research and Innovation, and the UKB. Sequencing was carried out in two centers (the deCODE facility in Reykjavik, Iceland, and the Wellcome Sanger Institute in Cambridge, UK). Genomic DNA underwent paired-end sequencing on Illumina NovaSeq6000 instruments with a read length of 2×151 and an average coverage of 32.5×^15,16^. Conversion of sequencing data in BCL format to FASTQ format and the assignments of paired-end sequence reads to samples were based on ten-base barcodes, using bcl2fastq (v.2.19.0). Initial QC was performed by deCODE and Wellcome Sanger, which included sex discordance, contamination, unresolved duplicate sequences, and discordance with microarray genotyping data checks.

UKB genomes were processed at AstraZeneca using the provided CRAM format files. A custom-built Amazon Web Services cloud compute platform running Illumina DRAGEN Bio-IT Platform Germline Pipeline (v.3.7.8) was used to align the reads to the GRCh38 genome reference and to call small variants. Variants were annotated using SnpEff (v.4.3)^17^ against Ensembl (Build 38.92)^18^. 45,755 sequences remained after removing contaminated sequences (verifybamid_freemix ≥ 0.04) using VerifyBAMID^19^ or that had low CCDS coverage (<94.5% of CCDS r22 bases covered with ≥10-fold coverage).

### Genome-wide association studies

The contribution of rare variants to common disease has, until recently, only been assessed for a subset of complex traits. The gnomAD, which includes exome and genome sequencing data of 141,456 individuals, constitutes the largest publicly available next-generation sequencing resource to date^20^. While this resource has undeniably transformed our ability to interpret rare variants and discover disease-associated genes, it is unsuited to the systematic assessment of the contribution of rare variation to human disease, as it lacks linked phenotypic data. We took forward variants that passed the variant QC as described in Wang et al.^3^, which had an MAC > 5. We tested the 2,751,139 variants identified in at least six individuals from the 45,755 predominantly unrelated European ancestry UKB exomes. Variants were required to pass the following quality control criteria: minimum coverage 10X; percent of alternate reads in heterozygous variants ≥ 0.2; binomial test of alternate allele proportion departure from 50% in heterozygous state P > 1 × 10−6; genotype quality score (GQ) ≥ 20; Fisher’s strand bias score (FS) ≤ 200 (indels) ≤ 60 (SNVs); mapping quality score (MQ) ≥ 40; quality score (QUAL) ≥ 30; read position rank sum score (RPRS) ≥ −2; mapping quality rank sum score (MQRS) ≥ −8; DRAGEN variant status = PASS; variant site is not missing (that is, less than 10X coverage) in 10% or more of sequences; the variant did not fail any of the aforementioned quality control in5% or more of sequences; the variant site achieved tenfold coverage in30% or more of gnomAD exomes, and if the variant was observed in gnomAD exomes,50% or more of the time those variant calls passed the gnomAD quality control filters (gnomAD exome AC/AC_raw ≥ 50%). Variant-level P values were generated by adopting Fisher’s exact two-sided test. Three distinct genetic models were studied for binary traits: allelic (A versus B allele), dominant (AA + AB versus BB), and recessive (AA versus AB + BB), where A denotes the alternative allele, and B denotes the reference allele. For quantitative traits, we adopted a linear regression (correcting for age, sex, and age × sex) and replaced the allelic model with a genotypic (AA versus AB versus BB) test. For ExWAS analysis, we used a significance cut-off of P ≤ 1 × 10−8. To support the use of this threshold in this study, we performed an n-of-1 permutation on the binary and quantitative trait dominant model ExWAS.

### Gene collapsing analyses

To perform collapsing analyses, we aggregate variants within each gene that fit a given set of criteria, identified as qualifying variants^21^. Overall, we performed 11 non-synonymous collapsing analyses, including 10 dominant and one recessive model, plus an additional synonymous variant model as an empirical negative control. In each model, for each gene, the proportion of cases was compared to the proportion of controls among individuals carrying one or more qualifying variants in that gene. The exception is the recessive model, where a participant must have two qualifying alleles, either in homozygous or potential compound heterozygous form. Hemizygous genotypes for the X chromosome were also qualified for the recessive model. These models were designed to collectively capture a wide range of genetic architectures. They vary in terms of allele frequency (from private up to a maximum of 5%), predicted consequence (for example, PTV or missense), and REVEL and MTR scores. Based on SnpEff annotations, we defined synonymous variants as those annotated as ‘synonymous_variant’. We defined PTVs as variants annotated as exon_loss_variant, frameshift_variant, start_lost, stop_gained, stop_lost, splice_acceptor_variant, splice_donor_variant, gene_fusion, bidirectional_gene_fusion, rare_amino_acid_variant, and transcript_ablation. We defined missense as: missense_variant_ splice_region_variant, and missense_variant. Non-synonymous variants included: exon_loss_variant, frameshift_variant, start_lost, stop_gained, stop_lost, splice_acceptor_variant, splice_donor_variant, gene_fusion, bidirectional_gene_fusion, rare_amino_acid_variant, transcript_ablation, conservative_inframe_deletion, conservative_inframe_insertion, disruptive_inframe_insertion, disruptive_inframe_deletion, missense_ variant_splice_region_variant, missense_variant, and protein_altering_variant.

Collapsing analysis P values were generated by using a Fisher’s exact two-sided test. For quantitative traits, we used a linear regression, correcting for age, sex.

For all models, we applied the following quality control filters: minimum coverage 10X; annotation in CCDS transcripts (release 22; approximately 34 Mb); at most 80% alternate reads in homozygous genotypes; percent of alternate reads in heterozygous variants≥0.25 and≤0.8; binomial test of alternate allele propor tion departure from 50% in heterozygous state P > 1 × 10−6; GQ ≥ 20; FS ≤ 200 (indels) ≤ 60 (SNVs); MQ ≥ 40; QUAL ≥ 30; read position rank sum score≥−2; MQRS≥−8; DRAGEN variant status=PASS; the variant site achieved tenfold coverage in ≥ 25% of gnomAD exomes, and if the variant was observed in gnomAD exomes, the variant achieved exome z-score ≥ −2.0 and exome MQ ≥ 30.

To quantify how well a protein-coding gene is represented across all individuals by the exome sequence data, we estimated informativeness statistics for each studied gene on the basis of sequencing coverage across the available exomes.

### Burden heritability and burden genetic correlation

GWAS have identified hundreds of independent loci associated with IDPs, and the heritability explained by common variants across IDPs has also been estimated. However, the contribution of rare coding variants to heritability is unclear. We used BHR^22^ (https://github.com/ajaynadig/bhr) to estimate the heritability explained by the gene-wise burden of rare coding variants across sleep traits. As recommended in BHR, the variants were stratified into bins according to allele frequency and functional categories. Ultrarare was defined as MAF < 1 × 10−5, and rare was defined as 1 × 10−5 ≤ MAF < 1 × 10−3. Functions of the variants were defined in the variant annotation section. All BHR analyses were run with the baseline file provided by ref^22^. The variant-level summary statistics derived from SAIGE-GENE+ outputs were used, and effect sizes of quantitative traits were obtained directly from SAIGE-GENE+ outputs, while effect sizes of binary traits were calculated using allele frequency among cases and controls.

Univariate BHR analysis was performed to estimate the burden h2 of each sleep trait, and we aggregated ultrarare and rare burden heritability together as total burden heritability and compared it with the common-variant heritability of each trait, which was obtained using linkage disequilibrium score regression^23^. Bivariate BHR analysis was then used to compute the genetic correlation rg. The burden genetic correlation between LoF and missense variants within each IDP was estimated. Burden genetic correlations across IDPs using ultrarare and rare LoF variants were also obtained.

### Conditional analysis adjusting for nearby common variation

To assess whether the detected rare variant signals were independent of nearby common variation, we repeated the gene-based collapsing analyses for the significant genes while adjusting for common variants in their surrounding regions. We first performed association testing of common variants (MAF > 0.5%) within a ±500-kb window of each significant gene using PLINK v2.0 (https://www.cog-genomics.org/plink/2.0/) and UK Biobank v3 imputed genotype data. We then clumped the association results using thresholds of P < 1 × 10⁻⁵ and r² < 0.01. The resulting set of clumped common variants was included as covariates when rerunning the gene-based collapsing analyses for these genes.

### Functional enrichment analysis and tissue expression

GO was selected as the gene set database to perform the enrichment analysis. GO terms were categorized as Biological Process, Cellular Component, and Molecular Function. To gain more insight into how genes may influence IDPs, we examined the enrichment of significant genes in 30 general tissues and 54 specific tissues from GTEx^24^ using FUMA^25^.

